# A cumulative inertia phenomenon explains anomalous long-distance transport in eukaryotic cells

**DOI:** 10.1101/339390

**Authors:** Sergei Fedotov, Nickolay Korabel, Thomas A. Waigh, Daniel Han, Victoria J. Allan

## Abstract

We demonstrate the phenomenon of cumulative inertia in intracellular transport involving multiple motor proteins in human epithelial cells by measuring the empirical survival probability of cargoes on the microtubule and their detachment rates. We found the longer a cargo moves along a microtubule, the less likely it detaches from it. As a result, the movement of cargoes is non-Markovian and involves a memory. We observe memory effects on the scale of up to 2 seconds. We provide a theoretical link between the measured detachment rate and the super-diffusive Levy walk-like cargo movement.

---

Intracellular transport of cargoes along microtubules is a classical example of active transport [1, 2]. It is critical to cellular function and it is a challenging statistical problem from the viewpoint of active matter physics [3–7]. *In-vitro* experiments show that the distance travelled by cargoes substantially increases when the cargoes are transported by multiple motor proteins [8]. Various models have been developed that aim to explain how motors achieve long range transport along microtubules [9–20].

In this Letter we reveal a new mechanism for anomalous directional persistence of cargoes in human cells: *the phenomenon of cumulative inertia*. Experimentally we found the longer a cargo moves along a microtubule, the less likely it will detach from it. To our knowledge our data provides the first direct measurement of the mesoscopic detachment rate as a decreasing function of the running time. We found that this time follows a heavy tailed Pareto distribution which leads to a Levy walklike movement of cargoes. Since the observed detachment probability depends on how long the cargo has been moving, this active transport involves memory and it exhibits a typical non-Markovian behaviour. In contrast, for systems with no memory, the mesoscopic detachment rate will not depend on the running time. The effect of cumulative inertia is well studied in social and behavioural sciences [21]. It has been found that the longer individuals remain in a state (a job, a residence, a state of mind, etc.), the less likely they will change this state in the future [22]. “Inertia” in these cases is accumulated over the duration of employment, residence time, etc. which follows a Pareto distribution. In recent years it has become a renewed area of interest in studies of human behaviour involving heavy tailed or Pareto distributions for activity times [23], such as sending e-mails, web surfing, etc. Our *in vivo* experiments show that cargo movement resemble the pattern of non-Poisson human activities: cargo transport includes rapid jiggling events separated by long periods of ballistic movements.

Recently, it has been discovered that active cargo transport *in-vivo* self-organizes into Levy walks [24]. Levy walks describe a wide spectrum of biological processes, such as T-cells migrating in brain tissue, collective behaviour of swarming bacteria and animals optimizing their search for sparse food [25]. Endosomal Levy dynamics involves long flights in one direction due to the active movement along microtubules driven by multiple motors. When all active motors disengage, the cargo complexes detach from the microtubule and reattach to a new microtubule heading in another direction [24]. The travel distances have power-law distributions with diverging variances [25–29] that explains the anomalously long flights of cargo complex. In Ref. [24] the authors proposed the concept of memoryless self-reinforced directionality to demonstrate the emergence of Levy walks.

However, the unanswered question remains: what is the precise mesoscopic kinetic mechanism of anomalous directional persistence? To answer this question, we performed *in vivo* experiments recording thousands of trajectories of intracellular lipid bound vesicles in live retinal pigment epithelium (RPE) cells and human bone osteosarcoma epithelial (U2OS) cells. We found similar results for both cell lines. Therefore, we report only results for RPE cells since a microscope with higher resolution was used to image them (see SI for experimental details). In Fig. 1, we illustrate Levy-like trajectories of vesicles inside RPE cells which consist of long persistent runs in one direction separated by rapid jiggling events when vesicles change direction.

**FIG. 1:**
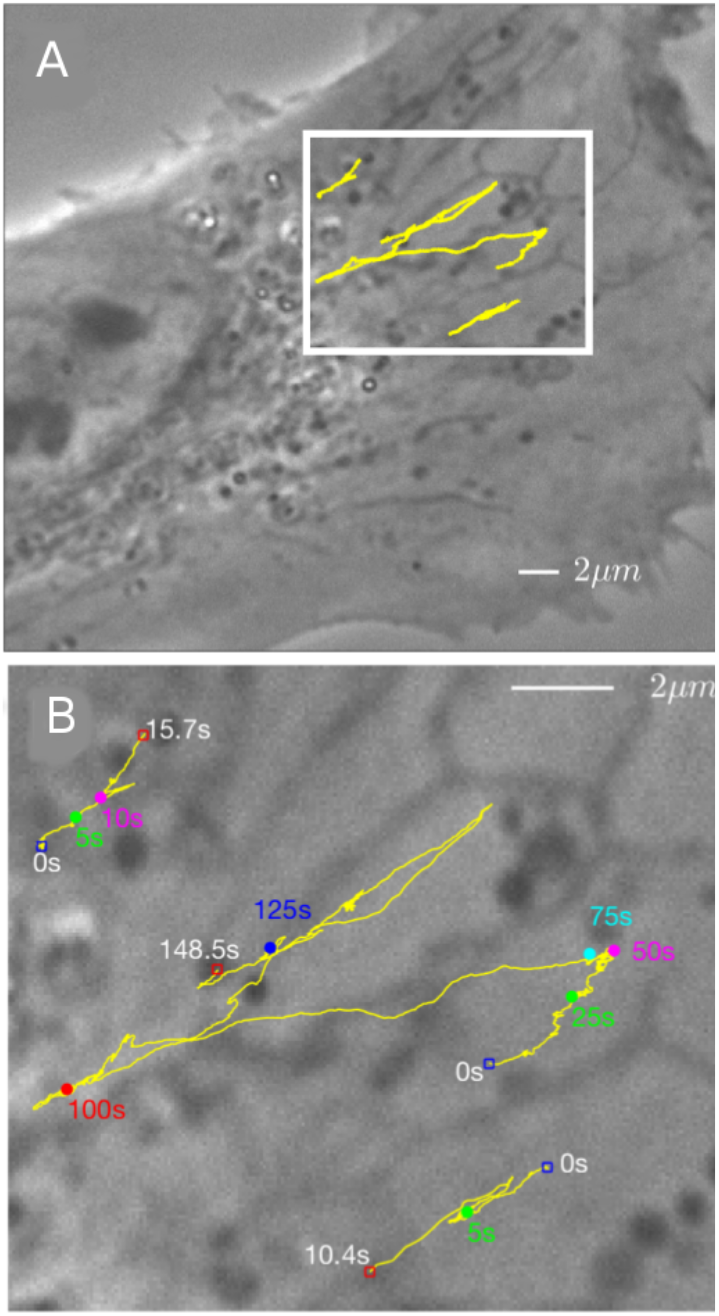
Cargoes movement along microtubules visualized in a living RPE cell (panel A). Panel B shows an enlargement of the boxed region in A. Trajectories (yellow) consists of long flights in one direction and multiple turnings. Numbers indicate the time progress. The longest trajectory (148.5 s) in the panel B consists of 900 flights (see SI for the description of the segmentation procedure).

One of our aims is to measure two important statistical characteristics of cargo transport: the empirical survival probability of cargoes on the microtubule and the empirical detachment rate. We estimated the survival probability by using the non-parametric Kaplan-Meier estimator [30]. We found a good agreement between the empirical survival probability Ψ and the heavy tailed Pareto distribution:

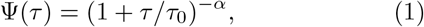

with the anomalous exponent *α* = 1.6 ± 0.17 and *τ*_0_ = 0.17 ± 0.1 s up to 1 – 2 seconds (Fig. 2). The empirical detachment rate is also found to be a decreasing function of the flight time *τ* (inset Fig. 2). This means that the longer a cargo remains on a microtubule, the less likely it will detach from it (*the phenomenon of cumulative inertia*). The empirical detachment rate has a good fit with the rate inversely proportional to the flight time *τ*:

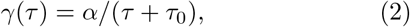

with the same anomalous exponent *α* = 1.6 ± 0.17 and *τ*_0_ = 0.17±0.1 s. Note the relationship between Ψ(*τ*) and 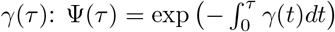. The time dependent detachment rate *γ*(*τ*) has the following meaning: the product *γ*(*τ*)Δ*τ* defines the conditional probability of cargo detachment in the interval (*τ, τ* + Δ*τ*) given that it has moved along the microtubule in the time interval (0, *τ*). For memoryless cargoes this rate will be constant and will not depend on how long the cargo has moved before. It is well-known that the empirical rate is notoriously difficult to estimate [30] since it contains the derivative of the empirical survival function. Therefore, the empirical survival probability (which is an integral quantity) has a smoother behaviour compared to the empirical rate function (Fig. 2).

**FIG. 2:**
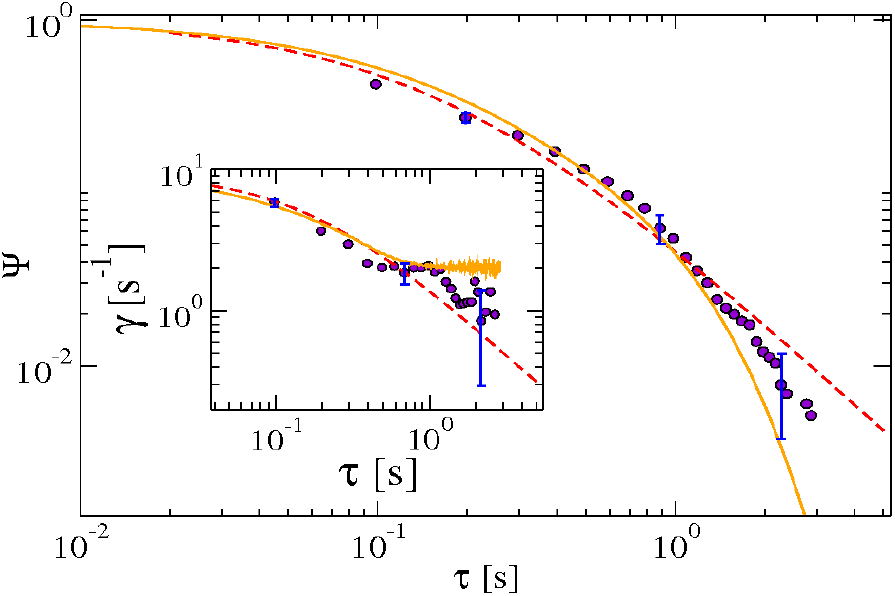
Experimentally determined survival function Ψ(*τ*) (dots) as a function of time *τ* decays as a power law on intermediate time scale with anomalous exponent *α* = 1.6 ± 0.17 and a time scale parameter *τ*_0_ = 0.17 ± 0.1 s. The red dashed curve is a power law fit Eq. (1) with the same α and *τ*_0_. The orange solid curve is the survival function obtained in numerical simulations of the cargo dynamics with 6 kinesin and 6 dynein motors. Details of simulations are given in the SI sec. IV. Inset: corresponding experimental empirical rate function *γ*(*τ*) (dots) as a function of time is inversely proportional to *τ*, Eq. (2), on intermediate time scale (red dashed curve) with the same *α* and *τ*_0_ as in the main figure. The orange solid curve is the rate function obtained in numerical simulations.

The value of experimental exponent *α* = 1.6 ± 0.17 falls in the interval 1 < *α* < 2. This is an extraordinary finding since it shows that for this interval the survival function has finite mean, 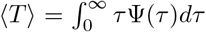, but divergent second moment [25]. Specifically the lack of the second moment leads to the emergence of the Levy walklike trajectories of vesicles (Fig. 1). Such trajectories exhibit sub-ballistic super-diffusive behaviour. We also obtain a good power law fit with the anomalous exponent (*α* ≃ 1.5 ± 0.17) for the probability density of flight lengths (Fig. 3) which confirms the Levy walk-like nature of the vesicles motion. The velocity v of each flight was assumed to be constant. The probability density of flight length is obtained from the survival function as:

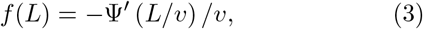

where Ψ′(*z*) = *d*Ψ(*z*)/*dz*. In Fig. 3 the distribution of flight velocities is also shown.

**FIG. 3:**
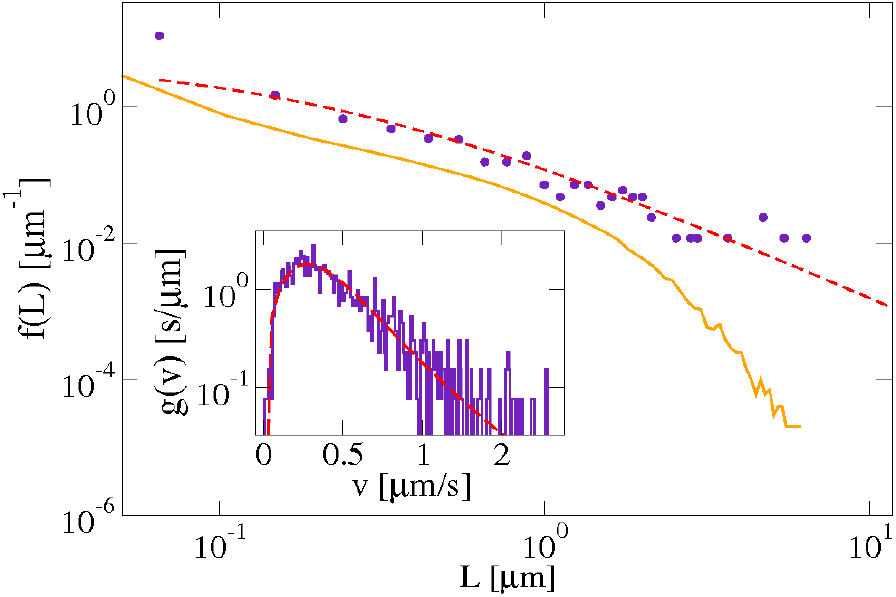
Main panel: The experimental flight length probability density *f*(*L*) (blue dots) fitted with Eq. (3) (red dashed curve) with *α* = 1.5 ± 0.17, *τ*_0_ = 0.33 ± 0.1 s and the average velocity of the cargo *v* = 0.8 *μ*m/s. The flight length distribution obtained in numerical simulations of the cargo dynamics with 6 dynein and 6 kinesin motors (orange solid curve) decays exponentially for *L* > 1 μm. Parameters of the simulations are given in SI. The inset shows the distribution of experimental flight velocities *g*(*v*) (blue solid curve) approximated with the Burr density (red dashed curve) with parameters 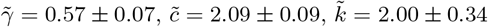. The average velocity of the cargo is *v* = 0.8 *μ*m/s.

Another important quantitative measure of cumulative inertia is the mean residual time 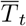 given that the cargo has survived on the microtubule up to time *t. In vivo* experiments we found that 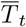 increases linearly in time *t* already travelled, see Fig. 4. The longer the cargo remains on the microtubule, the larger the mean residual time, so the inertia is accumulated. This behaviour is drastically different to memoryless systems where 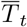 is constant and does not depend on the prehistory. The data can be well explained by the conditional survival function for a random attachment time time time *T*, Ψ_*c*_ (*t, τ*) = Pr{*T* > *t* + *τ*|*T* > *t*}. In our case:

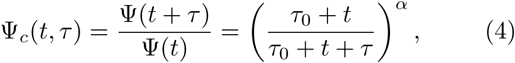

is an increasing function of time *t* already spent on the microtubule for a fixed *τ*. The behaviour of Ψ_*c*_ (*t, τ*) is illustrated in the inset of Fig. 4. The mean residual time 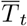 can be obtain as [31]:

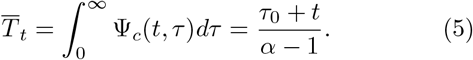

**FIG. 4:**
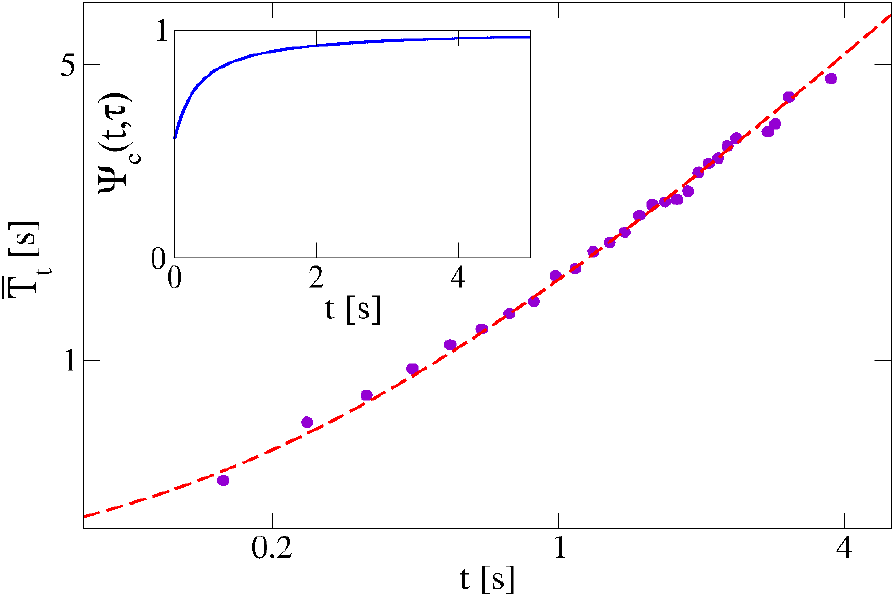
Main panel: The experimental mean residual time 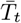 of flights (blue dots) linearly increases (notice logarithmic scales on both the horizontal and vertical axes) with the running time *τ*. The theoretical prediction Eq. (5) with parameters *τ*_0_ = 0.24 ± 0.1 s and *α* = 1.8 ± 0.17 (red dashed curve) is in good agreement with the experiment. Inset: Illustration of the increasing conditional survival function Ψ_*c*_(*t, τ*) given by Eq. (4) with *τ*_0_ = 0.2 s, *α* = 1.8 and *τ* = 0.1 s.

We found a good agreement between experimental mean residual time 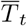 and Eq. (5) with *τ*_0_ = 0.24 ± 0.1 and the anomalous exponent α = 1.8±0.17 [32], Fig. 4. This cumulative inertia with 1 < *α* < 2 explains the dramatic increases of the travelled distance typical for Levy walk. Since this is a non-Markovian effect with memory, our explanation of anomalous long distance transport is completely different from the idea of memoryless self-reinforced directionality [24] when the probability *P*(*L*) of traveling in some direction grows with the distance *L* already travelled. This probability can be obtained in terms of the conditional survival function Ψ_*c*_(*t, τ*) as *P*(*L*) = Ψ_*c*_(*L*/*v, τ*).

Our next aim is to provide a theoretical link between the empirical detachment rate and the experimental Levy walk-like trajectories. Define the probability density function *ξ*(*t*, x, *φ, τ*) which gives the probability to find a cargo at point x = (*x, y*) at time t that moves with the velocity *v* in the direction ***θ*** = (cos *φ*, sin *φ*) and having started the move a time *τ* ago. Here *φ* is the angle between the direction of movement ***θ*** and the *x* axis. We assume that as long as the cargo detaches from the microtubule it immediately reattaches to another microtubule and thereby changes the direction of movement. The governing equation for *ξ*(*t*, x, *φ, τ*) takes the form [33]

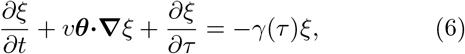

with the boundary condition

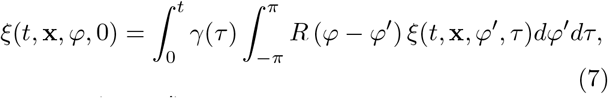

where *R*(*φ − φ*′) is the probability density of the reorientation from *φ*′ to *φ* such that 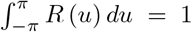. In Ref. [24] the flights were statistically isotropic with *R*(*u*) = 1/2*π*. In our experiments we observe quasi onedimensional trajectories (Fig. 1) and therefore use the following distribution *R*(*u*) = *δ*(*u*)/2 + *δ*(*u* − *π*)/2 [34]. The effective detachment rate Eq. (2) with the anomalous exponent 1 < *α* < 2 applied in Eq. (6) allows us to explain the emergence of Levy walk as a result of anomalous cumulative inertia phenomena. We obtain the mean squared displacement (msd) which exhibits sub-ballistic super-diffusive behaviour 〈x^2^(*t*)) ≈ *t*^3−α^ (SI sec. IV). For Levy walk with 1 < *α* < 2 the ensemble and time averaged msds differ only by a factor of 1/(*α* − 1) [35–37]. The msd calculated along single trajectories (corrected for a non zero drift and averaged over all trajectories) was found to grow as *t*^3−α^ with the anomalous exponent *α* ≃ 1.5 ± 0.17 consistent with the behaviour of the rate and survival functions (SI sec. III).

The question arises: what is the microscopic mechanism of the decreasing effective detachment rate *γ*(*τ*)? A first insight could be obtained from a classical Klumpp-Lipowsky model [9]. However, we note that this model could not explain the power law Pareto distribution for the survival probability. Consider the cargo which is pulled by multiple motors. We assume that initially the cargo attaches to the microtubule with a single motor. As the cargo moves along the microtubule, the number of engaged motors *N*(*t*) varies from 1 to 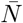. We define the random detachment time T as the time when all active motors together with the cargo detach from the microtubule. In order to obtain the effective detachment rate *γ*(*τ*), one can define the survival function Ψ(*τ*) of the cargo to remain on the microtubule as the probability Ψ(*τ*) = Pr{*T > τ*} = 1 − Pr{*N*(*τ*) = 0|*N*(0) = 1} [9]. The detachment rate is then [38] (SI sec. II):

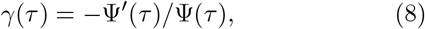

Note that this decreasing time-dependent detachment rate γ is essentially different from the effective constant unbinding rate obtained in Ref. [9] from equilibrium conditions. Motors attach to the microtubule and detach from it with rates *π_n_* and *ϵ_n_* (*n* is the number of engaged motors). To understand intuitively the reason why the effective rate *γ*(*τ*) decreases with the flight time, consider the first event of attachment of cargo complex with one motor. Initially the rate *γ*(0) = *ϵ*_1_. If the second motor attaches before the first motor detaches, the load is shared between two motors and the detachment rate *ϵ*_2_ decreases, *β*_2_ < *ϵ*_1_. As a result of this stochastic dynamics, the number of participating motors increases and therefore the detachment probability of the cargo decreases with the flight time. Cumulative inertia occurs due to multiple attachment and reattachment of motors before the cargo finally detaches from the microtubule. This leads to a dramatic increase of the travelled distance due to the directional persistence [8]. Our numerical results support this idea and show a decreasing effective detachment rate *γ* (inset Fig. 2). The empirical survival function can be approximated with the survival function corresponding to the Klumpp-Lipowsky model (Fig. 2).

*In vitro* experimental data indicates that adding just one extra motor increases cargo run lengths by at least one order of magnitude compared to the distance traveled by a single motor [8]. In what follows we consider the cargo pulled by two motors and show how the essential improvement of travel distance occurs. To obtain the rate *γ* defined in Eq. (8), we calculate the survival function (see SI sec. II):

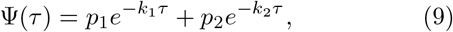

where 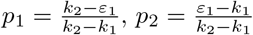. Here *k*_1_ and *k*_2_ (*k*_1_ < *k*_2_) are the solution of the quadratic equation *k*^2^ − (*π*_1_ + *ε*_1_ + *ε*_2_)*k* + *ε*_1_*ε*_2_ = 0. If *ε*_1_ < *k*_1_ (which is a typical condition for conventional motors), then the rate γ(τ) is *always* a decreasing function of the running time. The detachment rate *γ*(*τ*) takes the maximum value at *τ* = 0, *γ*(0) = *p*_1_*k*_1_ + *p*_2_*k*_2_ = *ε*_1_. In the long time limit, the detachment rate *γ*(*τ*) decreases and tends to the constant value *k*_1_, such that *k*_1_ < *ε*1. This explains the dramatic increase of running length of a cargo with two motors and the non-Markovian nature of cargo movement.

The survival function Eq. (9) has an interesting biological interpretation. The cargo movement can be viewed as one that involves a mixed population of two motors with different properties. If we extend this idea for a heterogeneous population of motors for which the rates k are gamma distributed with the probability density function *f*(*k*) = *τ*_0_*k*^*α*−1^*e*^−*τ*_0_*k*^/Γ(*α*), the effective survival function takes the form 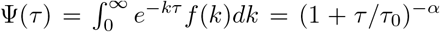 consistent with Eq. (1).

In conclusion, we studied experimentally the highly non-Markovian anomalous multi-motor intracellular transport of cargoes inside cells. Directly measuring the mesoscopic detachment rate of cargoes from microtubules, we have demonstrated that the origin of the anomalous non-Markovian behavior is the cumulative inertia phenomenon. We provided evidence for this phenomenon in both bone and retina epithelial cells, but it is expected to occur in all cell types that use dyneins and kinesins to transport cargoes along microtubules. Such intracellular transport impacts on a vast range of cellular processes and their diseased states e.g. motor neuron disease and cancer. Improved non-Markovian modelling will lead to more accurate quantitative analysis of the kinetics of a huge range of processes in cellular physiology.

Our model suggests a way to regulate the long distance cargo transport by controlling the effective detachment rate. One way this could be achieved is by recruiting more dynein molecules to the cargo. Recently, it has been shown that the adaptors that link dynein to cargo differ in their likelihood of binding either one or two dyneins per adaptor molecule [39, 40]. Cargoes with the 2-dynein adaptors move faster and farther than those with 1-dynein adaptors. We predict that high levels of 2-dynein adaptors would decrease both *α* and detachment rates, leading to a transition from Brownian transport to a Lévy walk regime.

## Acknowledgments

The authors acknowledge financial support from the EPSRC Grant No. EP/J019526/1.

